# Comparative Proteomic Profiling of Receptor Kinase Signaling Reveals Key Trafficking Components Enforcing Plant Stomatal Development

**DOI:** 10.1101/2025.07.20.665823

**Authors:** Pengfei Bai, Minh Huy Vu, Chiaki Komatsu, Ophelia Papoulas, Kazuo Ebine, Akira Nozawa, Tatsuya Sawasaki, Takashi Ueda, Edward M. Marcotte, Keiko U. Torii

## Abstract

Receptor kinases are pivotal for growth, development, and environmental response of plants. Yet, their regulatory mechanisms and spatial dynamics are still underexplored. The ERECTA-family receptor kinases coordinate diverse developmental processes, including stomatal development. To understand the proteomic landscape of the ERECTA-mediated signaling pathways, we here report comparative analyses of the ERECTA interactome and proximitome by epitope-tagged affinity-purification (ET-AP) and TurboID-based proximity labeling (TbID-PL) mass-spectrometry, respectively. While ET-AP successfully recovered receptor complex components (e.g., TOO MANY MOUTHS), TbID-PL effectively captured transient associations with the components of endosomal trafficking, i.e., clathrin-mediated endocytosis (CME) machinery. We further identify that specific subfamily members of phosphatidylinositol-binding clathrin assembly proteins (PICALMs) interact with and synergistically regulate ERECTA internalization. Mutations of these *PICALMs* impair ERECTA endocytosis and lead to excessive stomatal clustering by dampening the downstream signaling output. Taken together, our work provides a proteomic atlas of the ERECTA signaling network and suggests that timely removal of receptor kinase by the endocytosis machinery is essential for active signal transduction enforcing stomatal patterning.

## Introduction

Receptor kinases (RKs) constitute one of the largest gene families in plants, playing a fundamental role in orchestrating growth, development, environmental responses, and immunity (*1–6*). Among them, those with an extracellular leucine-rich repeat (LRR) domain comprise the largest subgroups, with well-studied members including a phytohormone brassinosteroid receptor BRASSINOSTEROID INSENSITIVE1 (BRI1) (*7*) and innate immunity receptor FLAGELLIN SENSING2 (FLS2) (*8*). The ERECTA family of three LRR-RKs, ERECTA (ER), ERECTA-LIKE1 (ERL1), and ERECTA-LIKE2 (ERL2) regulates multiple aspects of developmental processes, including epidermal patterning, organ morphogenesis, vascular differentiation, and meristem activity (*9, 10*). The functional redundancy and synergy among these receptors are evident from the severe developmental defects observed in *er erl1 erl2* triple mutants, which exhibit increased stomatal density and clustering, reduced plant stature, and arrested inflorescence (*11, 12*). The pleiotropic nature of ERECTA signaling is mediated by interactions with EPIDERMAL PATTERNING FACTORs (EPFs) and EPF-LIKEs (EPFLs), a diverse family of endogenously secreted peptides that enable spatiotemporal regulation of receptor signaling (*13–16*).

ERECTA fine-tunes stomatal development by balancing the proliferation and differentiation of stomatal initial cells, meristemoid mother cells (MMCs), and dividing precursor cells called meristemoids (*12, 17*). Studies in the past decade have revealed the signal transduction pathways mediated by ERECTA in a few developmental contexts (*10*). Specifically, EPF2 is secreted from early stomatal-lineage cells and binds ERECTA to inhibit excessive stomatal formation. STOMAGEN/EPFL9 acts as an antagonistic peptide and competes with EPF2 for receptor binding, promoting stomatal differentiation (*15, 18, 19*). This ensures a tightly regulated developmental balance. ERECTA requires its co-receptor TOO MANY MOUTHS (TMM) for selective ligand recognition (*20–22*). Unlike ERECTA, TMM lacks a cytoplasmic kinase domain and therefore does not directly participate in intracellular signal transduction. Instead, ligand-activated ERECTA interacts with co-receptors, including BRI1-ASSOCIATED KINASE1/SOMATIC EMBRYOGENESIS RECEPTOR KINASE3 (BAK1/SERK3) and related SERKs, which trans-phosphorylate and activate ERECTA (*23, 24*). Receptor-like cytoplasmic kinases (RLCKs), BRASSINOSTEROID-SIGNALING KINASE 1 (BSK1) and BSK2 also function downstream, likely serving as a signaling scaffold that bridges ERECTA-mediated signaling to the MAP kinase cascade via YODA (YDA) (*25*).

The swift turnover of activated receptors involves multiple layers of control, with receptor endocytosis playing a crucial role in modulating signaling intensity, duration, and spatial distribution (*26*). Plant RKs undergo both constitutive and ligand-induced endocytosis, with BRI1 and FLS2 serving as prominent examples (*27–30*). Accumulating evidence shows that both receptors are internalized via clathrin-mediated endocytosis (CME) (*28, 31*). Members of the ERECTA family also undergo ligand-dependent endocytosis: ERL1 is trafficked to multivesicular bodies for vacuolar degradation in response to EPF1, a process inhibited by the antagonist peptide STOMAGEN, while ERL2 likewise internalizes (*32, 33*). Despite these observations, the molecular machinery directing the endocytosis of ERECTA family receptors remains unknown. The discovery of the TPLATE complex as a plant-specific adaptor that directly interacts with ubiquitinated cargo suggests a mechanism for cargo selectivity (*34, 35*). Yet, how distinct classes of receptors, such as developmental RKs, immune or hormone receptors, are differentially recognized by the evolutionarily conserved adaptor proteins remains unresolved.

The interactome of LRR-RLKs has been extensively profiled, revealing an integrated network of extracellular receptor-ligand interactions (*36*). While proteomic studies have advanced our understanding of RK signaling, its regulatory mechanisms and dynamic signaling processes still remain a challenge. Conventional affinity purification-mass spectrometry has been instrumental in identifying RK interactors (*37, 38*). However, it has inherent limitations in detecting transient or weak interactions, leaving a significant portion of the RK intracellular network unresolved (*39*). TurboID-based proximity labeling (TbID-PL) has emerged as a powerful approach for mapping low-abundant and transient interactors with high spatial and temporal resolution (*40–43*). Here, we employ a parallel proteomic approach using epitope-tagged affinity-purification (ET-AP) and TbID-PL from the same biological samples to construct a comprehensive proteomic map of the ERECTA signaling network. In addition to the canonical ERECTA signaling components, we identify a suite of adaptor and accessory proteins linked to CME (*35*). Among these, we discover that PICALMs, key regulators of CME, associate with ERECTA and play a crucial role in receptor internalization and homeostasis. *picalm* mutants hierarchically impact stomatal patterning. Further genetic analyses suggest that impaired PICALM functions dampen the canonical ERECTA-YDA-MAPK output. Collectively, our study establishes a proteomic atlas of the ERECTA signaling network and highlights the significance of CME-mediated receptor dynamics in maintaining robust RLK signaling and proper stomatal development.

## Results

### Comparative proteomic profiling of ERECTA-associated proteins

To unravel the full proteomic landscape of ERECTA-mediated cellular receptor signaling, it is imperative to simultaneously profile both the ERECTA-interacting proteome (i.e., those proteins that tightly associate with ERECTA) and proximity proteomes (i.e., those proteins in the close vicinity of ERECTA receptor complexes). To achieve this goal, we performed ET-AP and TbID-PL from the same Arabidopsis seedling extracts (see Materials and Methods; Fig. 1A). For this purpose, we generated transgenic Arabidopsis seedlings expressing the *ERECTA-HA-TbID* fusion protein under the control of the native ERECTA promoter in the *erecta* null allele, *er-105*. The expression of an ERECTA-HA-TbID fusion protein fully rescued the characteristic inflorescence and stomatal patterning defects of *er-105*, validating its biological functionality (fig. S1). Notably, immunoblot analysis verified efficient biotinylation of proximal proteins in the TbID-PL experiments and successful isolation of the fusion protein (Fig. 1B and fig. S2).

**Fig. 1.**
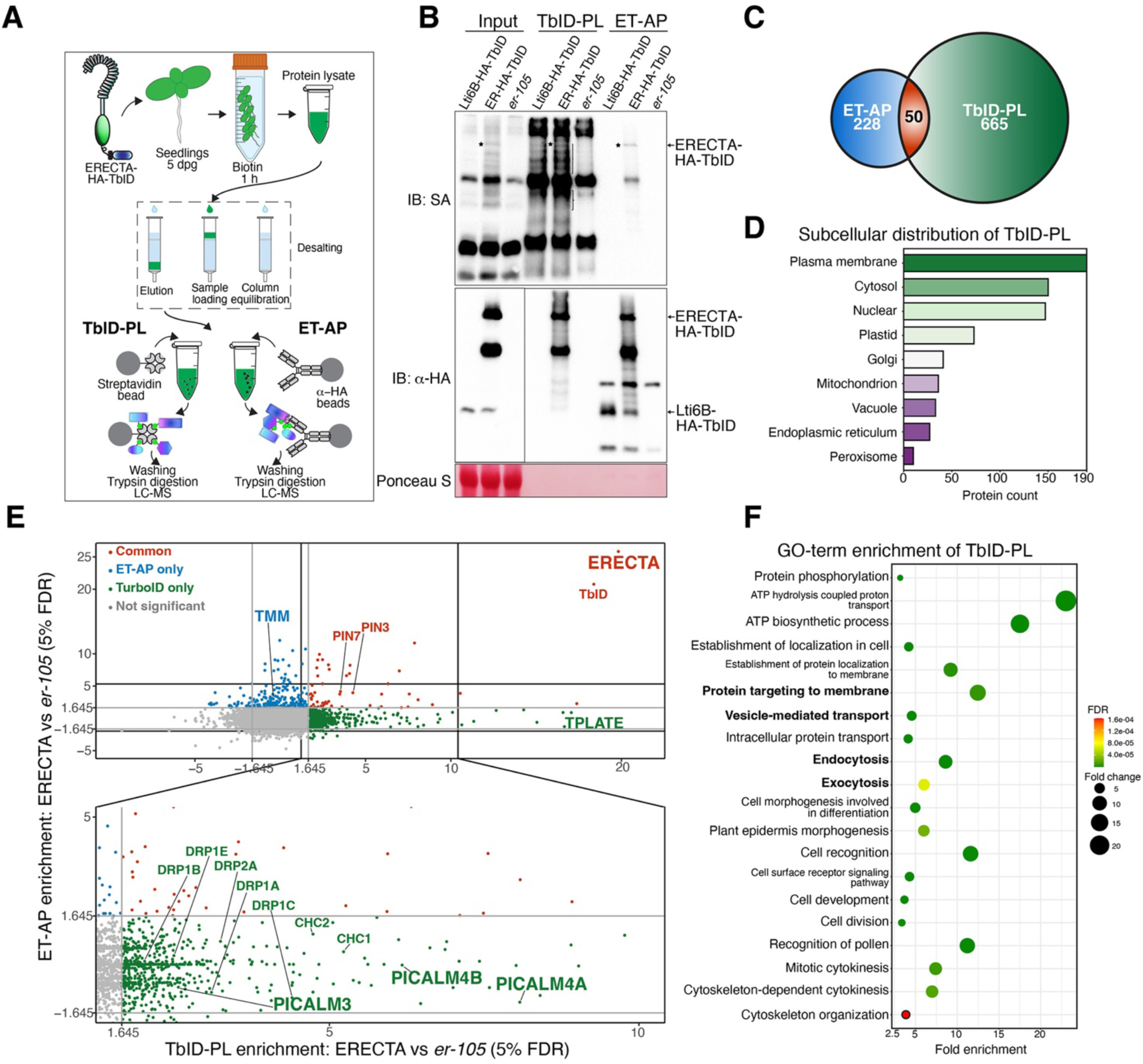
Comparative proteomic profiling of ERECTA-associated proteins. **(A)** Schematic representation of the experimental workflow for profiling ERECTA-associated proteins using TbID-PL and ET-AP. Parallel enrichments of biotinylated proteins and HA-tagged protein complexes were subjected to mass spectrometry analysis. **(B)** Immunoblot validation of ERECTA and its associated proteins. Streptavidin-HRP (SA-HRP) (top) confirms successful biotinylation of proximal proteins via TbID-PL. The α-HA immunoblot (middle) verifies the expression and enrichment of ERECTA-HA-TbID. Ponceau S staining (bottom) indicates loading control. **(C)** Venn diagram of identified proteins reveals 665 unique to TbID-PL, 228 unique to ET-AP, and 50 shared proteins. **(D)** Subcellular localization of TbID-PL enriched proteins, showing enrichment at the plasma membrane and cytosol, with additional representation in endomembrane compartments. **(E)** Scatterplot comparing protein enrichment between TbID-PL and ET-AP. Notable components of clathrin-mediated endocytosis (CME), such as PICALMs, DRPs, and TPLATE complex members, are preferentially captured by TbID-PL. In contrast, stable receptor partners like TMM are enriched via ET-AP. Some auxin transporters (PIN3, PIN7) are recovered in both methods. All TbID-PL and ET-AP experiments were performed using three biological replicates. Protein enrichment was quantified as log2-transformed ratios of normalized PSMs between experimental and control conditions and significance was assessed using a one-sided Z-test (z ≥ 1.645, BH-adjusted FDR). Proteins enriched in both approaches are shown in red, TbID-PL specific enrichments in green, and ET-AP specific in blue. **(F)** GO-term enrichment of TbID-PL-identified proteins reveals overrepresentation of membrane trafficking, endocytosis, protein localization, and epidermal development terms. Circle size represents fold enrichment; color indicates false discovery rate (FDR) values.

Through the parallel proteomic approaches, we identified 715 proteins via TbID-PL and 278 proteins via ET-AP, with 50 proteins shared between the two approaches (Fig. 1C and 1E, Data S1). We first examined the proteomes specifically enriched by each approach. TbID-PL prominently enriched transient and membrane-associated interactors, including proteins involved in CME, most notably phosphatidylinositol-binding clathrin assembly protein (PICALM) family members, the TPLATE complex, adaptor protein-2 (AP-2) complex and dynamin-related proteins (DRPs) (Fig. 1E, Data S1). On the other hand, ET-AP exclusively detected TOO MANY MOUTHS (TMM), a known regulator of stomatal patterning that forms a stable receptor complex with ERECTA (*22*) (Fig. 1E). It is important to note that TMM is an LRR-RLP with an extracellular and transmembrane region but lacks any recognizable cytoplasmic domain (*44*). Since TbID was fused to the cytoplasmic domain of ERECTA (Fig. 1A), the absence of TMM biotinylation by ERECTA TbID-PL indicates that the plasma membrane environment was preserved during labeling. Lastly, both approaches effectively captured ERECTA itself, along with shared interactors like auxin efflux transporters PIN3 and PIN7, hinting at a molecular link between ERECTA and auxin-mediated plant growth control (*45, 46*).

Consistent with our finding that TbID-PL preferentially captures dynamic and transient interactors of the RK ERECTA, the identified proteins are significantly enriched in Gene Ontology (GO) categories related to “protein targeting to membranes”, “vesicle-mediated transport”, and “intracellular protein transport and endocytosis” (Fig. 1F). Additional enriched categories include “plant epidermis morphogenesis”, “cell recognition”, “cell division and cytokinesis”, and “ATP hydrolysis and biosynthesis”, likely reflecting the broad range of biological processes mediated by the ERECTA signaling. Subcellular localization analysis further revealed that ERECTA-associated proteins are predominantly enriched at the plasma membrane and cytosol (Fig. 1D, Data S4), underscoring the spatial specificity of TbID-PL in detecting proximal proteins. Together, these findings highlight the complementary strengths of TbID-PL and ET-AP in defining the ERECTA-associated proteome. Specifically, our results indicate that TbID-PL exhibits higher sensitivity in detecting transient and dynamic interactors, whereas ET-AP identifies stable extracellular complexes, thereby providing a comprehensive view of ERECTA-associated proteomic landscape.

### Optimized TbID-PL enhances detection of ERECTA-signaling components and reveals compartment-specific proteomic profiles

Extensive genetic studies have identified the downstream components of ERECTA signal transduction pathways (*23, 25, 47–49*). However, we did not detect many such components through either ET-AP or TbID-PL, possibly due to their low abundance. It is increasingly recognized that TbID-PL in plants faces significant background noise owing to the endogenous biotin pools within plant cells (*50, 51*). To improve the detection of biologically-relevant proximitome while minimizing background noise, we optimized the labeling strategy by comparing the biotinylated proteomes of ERECTA-HA-TbID seedlings treated with exogenous biotin (with biotin, 1 hr) to a mock treated control (without biotin, 1 hr) (Fig. 2A and fig. S2C; see Materials and Methods). This approach effectively captured BAK1 (*23*), the transient co-receptor of ERECTA, as well as the downstream MAPKKK, YODA (*47*) (Fig. 2B, Data S2).

**Fig. 2.**
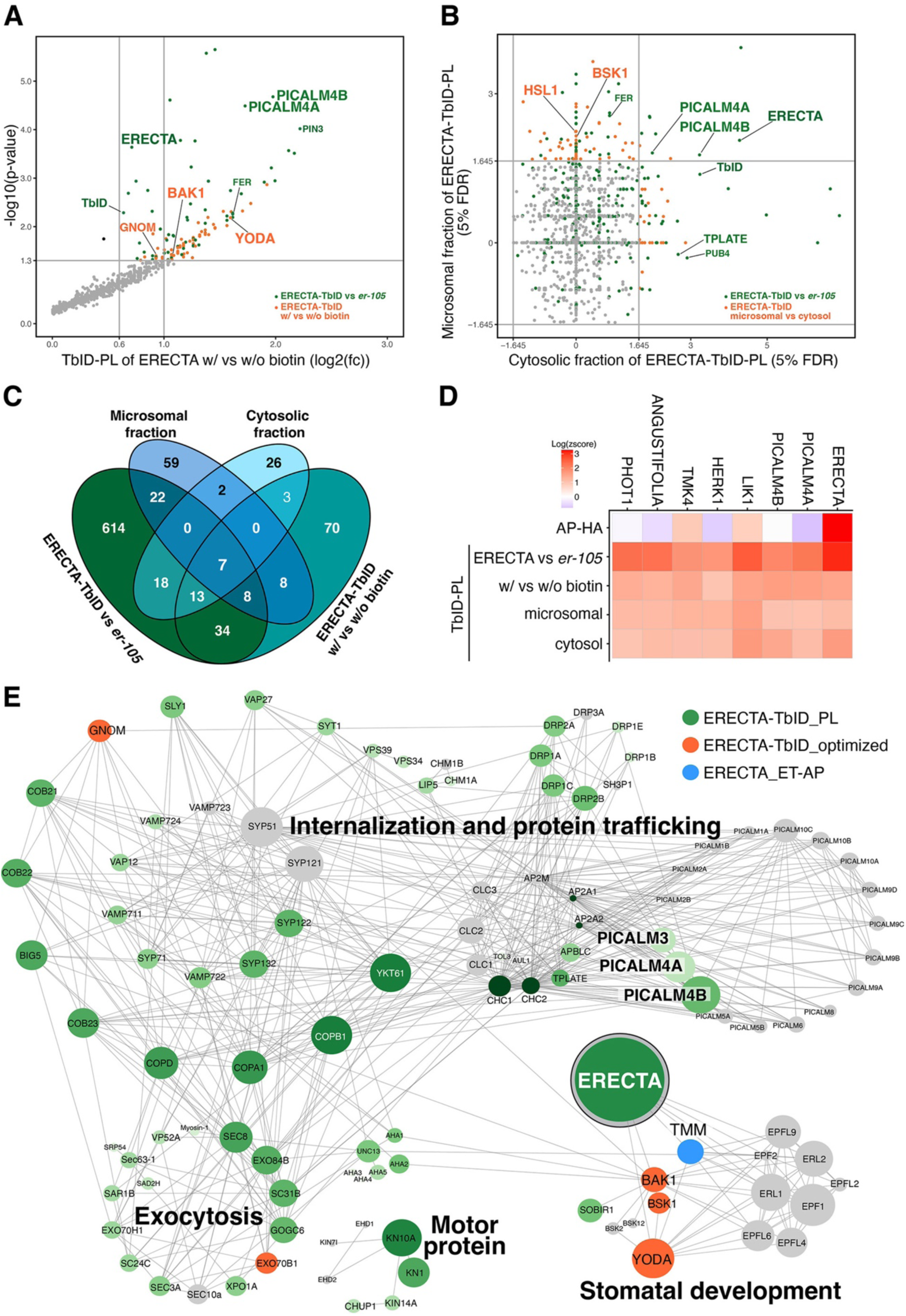
Optimized TbID-PL enhances detection of ERECTA-associated signaling components and compartment-specific proteomes. **(A)** Scatterplot showing differential enrichment of biotinylated proteins in ERECTA-TbID-PL with biotin versus without biotin treatments. Several known ERECTA signaling components, including BAK1, YODA (MAPKKK4) are identified (orange) alongside clathrin-mediated endocytosis proteins (e.g., PICALM4A, PICALM4B) identified (green) in Fig. 1. TbID-PL experiments were performed with three biological replicates. **(B)** Scatterplot comparing enrichment between microsomal and cytosolic fractionation in TbID-PL proteomes. Membrane-attached receptor-like cytoplasmic kinase BSK1 is predominantly enriched in the microsomal fraction, while CME components (e.g., PICALM4A, 4B) and ERECTA are detected in both fractions. TbID-PL experiments were performed with three biological replicates. **(C)** Venn diagram showing overlap among TbID-PL datasets derived from comparisons of ERECTA-HA-TbID vs *er-105*, +biotin vs -biotin, and microsomal vs cytosolic fractions. Seven candidates are consistently detected across all comparisons. **(D)** Heatmap showing normalized enrichment (log2 z-scores) of the top 7 consistently identified ERECTA-proximal proteins across four TbID-PL conditions. These include ERECTA itself, PICALM4A/B, and kinases implicated in receptor signaling (LIK1, HERK1, TMK4), as well as ANGUSTIFOLIA and PHOT1. **(E)** STRING network analysis of ERECTA-associated proteins identified by TbID-PL (green), optimized TbID-PL (orange), or ET-AP (blue). Proteins are clustered into major functional categories: internalization/protein trafficking, exocytosis, motor proteins, and stomatal development. ERECTA, CME components (PICALMs, TPLATE, DRPs), and signaling partners (e.g., BAK1, YODA, TMM) are highlighted. Node size reflects interaction confidence.

To enhance the membrane proximitome signals, we next performed microsomal fractionations, which retained higher levels of full-length ERECTA with reduced biotinylation background signals (fig. S2D). Through the ERECTA membrane TbID-PL, we are able to detect the membrane-attached receptor-like cytoplasmic kinase BSK1, which likely relays the ERECTA signaling (*25*) (Fig. 2B, Data S3). We additionally detected the LRR-RK HAESA-LIKE1 (HSL1), which has been shown to be expressed in stomatal-lineage cells and to modulate stomatal development (*52*) (Fig. 2B). Notably, the endocytic adaptor proteins PICALM4A and PICALM4B are robustly and consistently detected in all four TbID-PL conditions. Additional consistent ERECTA TbID-PL interactors include several RKs, TMK4 (*53*), HERK1 (*54*), LIK1 (*55*), as well as ANGUSTIFOLIA, which is a component of the stomatal polarity complex (*56*), and the blue-light receptor PHOT1 (*57*) (Fig. 2C and 2D). Together, our multilayered strategies of TbID-PL procedures effectively capture ERECTA signaling components, enable detection of compartment-specific proximitomes, and establish the core sets of proteins in the proximity of ERECTA that could guide future explorations.

### ERECTA interacts with the PICALM family of clathrin-mediated endocytosis (CME) components

Our ERECTA TbID-PL highlights the CME components as transient interactors of ERECTA. While ERECTA-family RKs are known to undergo endocytosis, no specific components that recognize ERECTA as a cargo had been identified (*32, 33, 58*). To explore the subcellular regulatory dynamics of ERECTA, we first analyzed its association with the CME machinery by constructing a landscape of the ERECTA network using STRING database analysis (*59*) (see Methods). The network grouped ERECTA-associated proteins into distinct functional categories: internalization and protein trafficking, exocytosis, motor protein function, and stomatal development (Fig. 2E). Remarkably, none of the components other than the category of stomatal development had been shown to directly interact with ERECTA, while they are overwhelmingly detected through TbID-PL.

Among the newly identified ERECTA associatome, several CME-related components, notably PICALM3, PICALM4A, and PICALM4B, captured our attention for their highly selective enrichment (Figs. 1 to 3). Importantly, PICALMs are known to function as adaptor proteins for specific cargos but are not required for general endocytosis (*60*). To gain insight into their potential relevance in stomatal development, we re-analyzed single-cell transcriptomic data (*61*) and found that only a subset of PICALMs from Group 1 and Group 2 are highly expressed across stomatal lineage cells (Fig.3, A to C). Among them, PICALM4A showed the strongest and broadest expression spanning meristemoids, stomatal lineage ground cells (SLGCs), guard mother cells (GMCs), and guard cells (GCs), while PICALM1A, PICALM3, PICALM4B were also expressed to varying degrees. The expression pattern guided our initial biochemical and genetic analysis. We therefore postulated that PICALMs may represent the yet unidentified adaptors for ERECTA. To address this hypothesis, we first surveyed the protein-protein interactions of the ERECTA cytoplasmic domain (ERECTA-CD) and members of the PICALM family throughout their five evolutionary subgroups (*62*) (Fig. 3B), using *in vitro* translated proteins in the Amplified Luminescent Proximity Homogeneous Assay Screen (AlphaScreen assay) (*63*) (Fig. 3D; See Materials and Methods). Indeed, we detected the highest interactions (i.e., normalized binding activity) of ERECTA-CD with PICALM3 (Subgroup 1) and PICALM4A (Subgroup 2a) followed by PICALM1A (Subgroup 1), consistent with their expression patterns in stomatal lineage cells. Additionally, PICALM2A, PICALM4B, PICALM5A, and PICALM6 exhibited positive but weaker interactions with ERECTA-CD (Fig. 3D). We further performed co-immunoprecipitation (Co-IP) assays using transgenic Arabidopsis seedlings expressing both *ERECTA-FLAG* and *PICALM4A-RFP* under their native promoters and demonstrated that ERECTA interacts with PICALM4A *in vivo* (Fig. 3E). Our transient protoplast co-immunoprecipitation assays also validated the interaction of ERECTA with PICALM3, PICALM4A, and PICALM4B (fig. S3C). Combined, these findings establish PICALM family members of CME as *in vivo* interactors of ERECTA, thereby implicating them in the endocytosis of ERECTA.

**Fig. 3.**
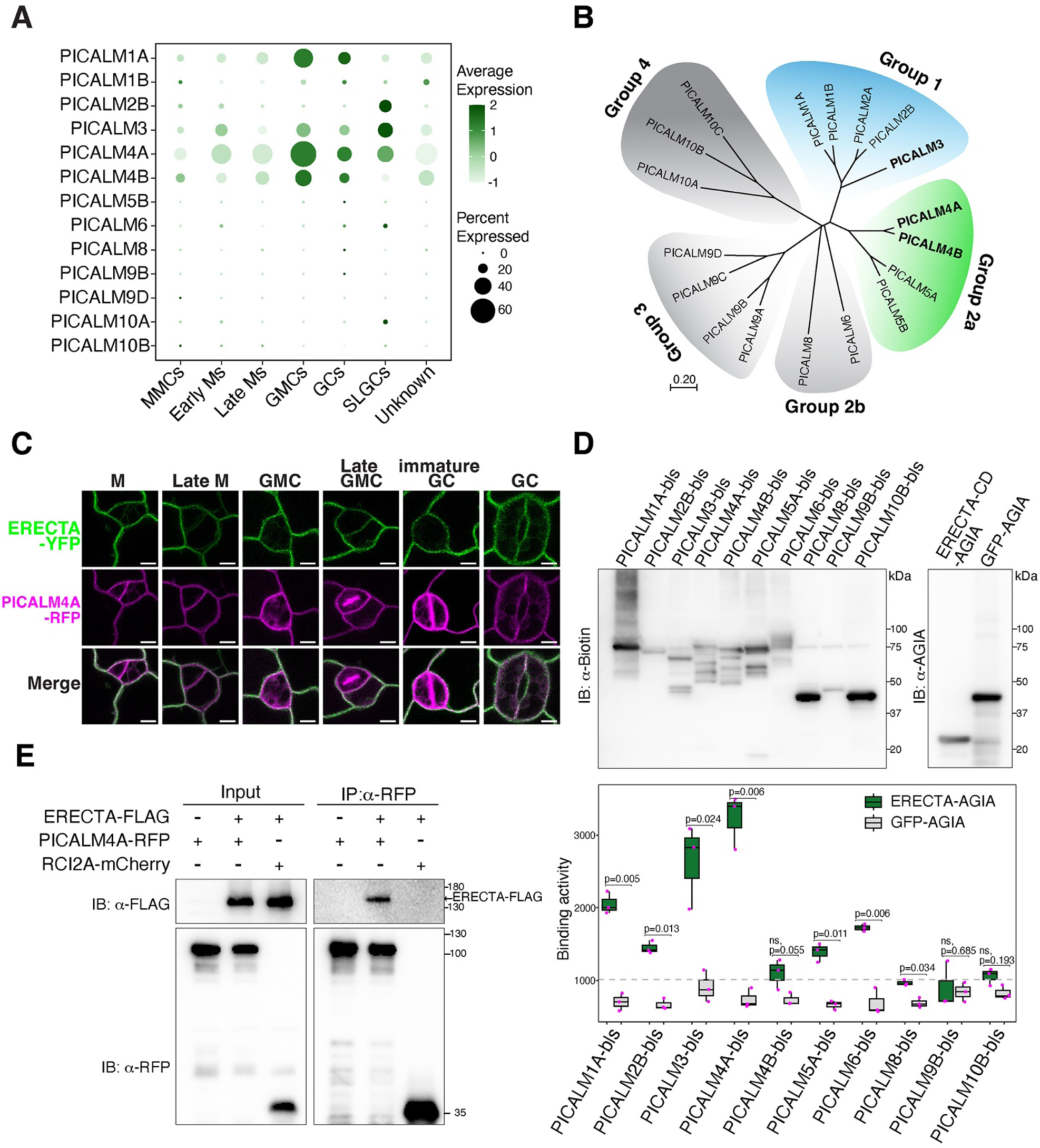
ERECTA selectively interacts with PICALM family of clathrin-mediated endocytosis (CME) components. **(A)** Bubble chart showing transcriptional expression of Arabidopsis PICALM family genes across stomatal lineage cell types from re-analyzed single-cell RNA-seq data. PICALM1, PICALM3, PICALM4A, and PICALM4B are present across different stages of stomatal lineage cells. PICALM1, PICALM4A, and PICALM4B are highly enriched in guard mother cell (GMC), guard cell (GC) stages, while PICALM2B and PICALM3 are highly enriched in stomatal lineage ground cells (SLGCs). The distinct expression of PICALMs suggests their potential roles in different stages of stomatal development. Dot size indicates the proportion of expressing cells; dot color reflects expression levels. **(B)** Phylogenetic tree of Arabidopsis PICALM family members. PICALM3 (Group 1, blue) and PICALM4A/B (Group 2a, green) cluster into distinct clades with high proximity to ERECTA based on TbID-PL. The phylogenetic tree illustrates evolutionary relationships and potential functional specialization within PICALM family. **(C)** Confocal microscopy showing subcellular colocalization of endogenous promoters-driven ERECTA-YFP and PICALM4A-RFP in cotyledon epidermal cells at multiple stomatal lineage stages. M, meristemoid; GMC, guard mother cell; GC, guard cell. Scale bars: 10 μm. ERECTA-YFP colocalizes with PICALM4A-RFP at the plasma membrane in stomatal lineage cells, except at cell plates for dividing cells. **(D)** *In vitro* protein-protein interacting assay between ERECTA and PICALM family proteins. ERECTA cytoplasmic domain and PICALMs were *in vitro* translated for the AlphaScreen assay. Quantification of normalized binding activity (right) confirms the preferential binding for PICALM3, PICALM4A, and PICALM1A. Boxplots represent mean with SD from three biological replicates and significance was assessed by unpaired t-test. **(E)** Co-immunoprecipitation assays validating interactions between ERECTA and PICALM proteins *in vivo*. Transgenic Arabidopsis seedlings expressing both ERECTA-FLAG and PICALM4A-RFP under their native promoters were subjected to α-RFP immunoprecipitation. For the original, uncropped gel blot images, see fig. S13.

### PICALMs colocalize with ERECTA and regulate its endocytosis

To elucidate the roles of PICALMs in ERECTA’s subcellular dynamics, we first examined colocalization of ERECTA with PICALM proteins and endocytic markers in stomatal lineage cells. Under mock conditions, ERECTA-YFP colocalized with PICALM4A at plasma membrane across stomatal lineage cells (Fig. 4A and 4B). Upon treatment with Brefeldin A (BFA), which inhibits ADP-ribosylation factor guanine-nucleotide exchange factor (ARF-GEF) activity and induces endosomal compartments (known as “BFA bodies”) (*64*), ERECTA-YFP accumulated within the BFA bodies in epidermal cells and strongly colocalized with PICALM4A-RFP, where PICALM4A-RFP signals appear to encapsulate ERECTA-YFP, suggesting PICALM-mediated recruitment of ERECTA into early endosomal structures (Fig. 4A, yellow arrowheads; 4E, Movie S1). To delineate the endocytic route of ERECTA, we next examined colocalization of ERECTA with PICALM4A upon wortmannin (Wm) treatment, which generates enlarged late endosomal compartments (Wm bodies, 0.5-1.0 µm in diameter) (*32*). ERECTA-YFP partially colocalized with PICALM4A in Wm bodies (Fig. 4B, yellow arrowheads; 4E), suggesting progression into late endosomes. ERECTA-YFP also strongly colocalized with the trans-Golgi network (TGN) marker SYP43-RFP and the early endosome marker ARA6-RFP at the plasma membrane under mock conditions. Following BFA treatment, ERECTA-YFP accumulated in endosomal compartments containing these markers, indicating its trafficking through the TGN and early endosomal pathways (Fig. 4C, yellow arrowheads; 4F). Additionally, ERECTA-YFP colocalized with the late endosome marker ARA7-RFP under Wm-treated conditions (Fig. 4D, yellow arrowheads; 4G), suggesting its progression through late endosomal compartments. In contrast, its association with the vacuolar membrane marker SYP22-RFP was limited (Fig. 4D, 4G). These results demonstrate that PICALM-mediated CME facilitates the dynamic trafficking of ERECTA-YFP from plasma membrane through the TGN and early-to-late endosomal compartments.

**Fig. 4.**
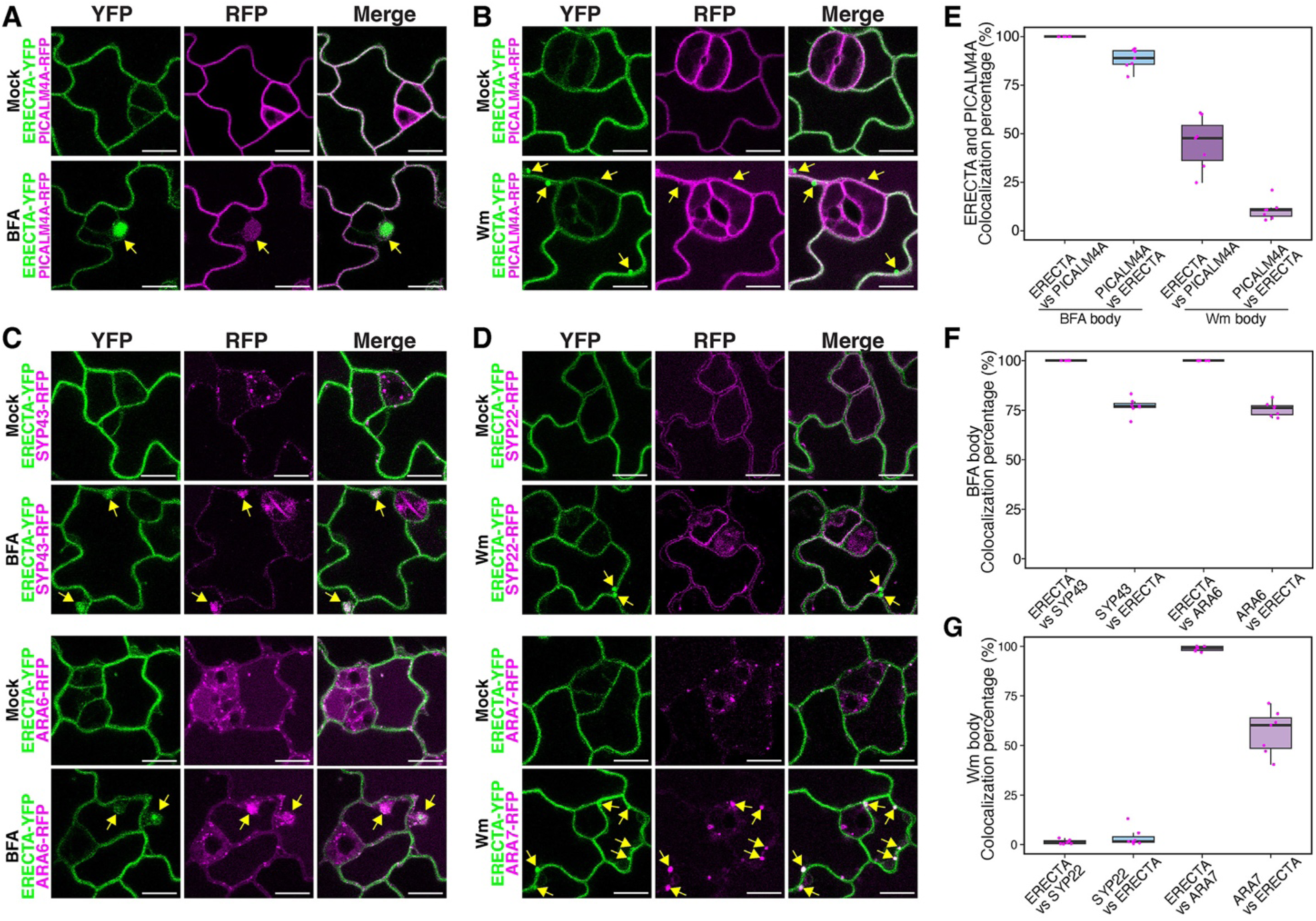
ERECTA-YFP colocalizes with PICALM4A and endosomal markers in stomatal-lineage cells. **(A)** Confocal microscopy showing colocalization of ERECTA-YFP with PICALM4A-RFP under mock and Brefeldin A (BFA)-treated conditions. BFA treatment results in BFA-induced endosomal compartments (BFA bodies, yellow arrows), where PICALM4A-RFP (magenta) encapsulates and colocalizes with ERECTA-YFP (green). **(B)** Confocal microscopy showing partial colocalization of ERECTA-YFP with PICALM4A-RFP under mock and wortmannin (Wm) treatments. Wm induces enlarged late endosomal compartments (Wm bodies, 0.5-1.0 µm in diameter; yellow arrows), in which ERECTA-YFP partially colocalizes with PICALM4A-RFP. **(C)** Colocalization of ERECTA-YFP with the trans-Golgi network (TGN) marker SYP43-RFP (magenta; top) and the plasma membrane/early endosome marker ARA6-RFP (magenta; bottom). Following BFA treatment, ERECTA-YFP (green) strongly colocalizes with both SYP43-RFP and ARA6-RFP in BFA bodies. **(D)** Confocal microscopy showing partial colocalization of ERECTA-YFP with the late endosome marker ARA7-RFP (magenta; bottom) and the vacuolar membrane marker SYP22-RFP (magenta; top). Minimal colocalization is observed under mock conditions. Upon Wm treatment, ERECTA-YFP colocalizes with ARA7-RFP in these enlarged endosomes. **(E-G)** Quantification of ERECTA-YFP colocalization with PICALM4A-RFP (E), SYP43 and ARA6 markers (F), SYP22 and ARA7 markers (G) in BFA bodies or Wm bodies. Colocalization percentage was calculated as the proportion of overlapping endosomes between the two proteins to the total endosome number of former protein. Boxplots represent median values with interquartile ranges. n = 7 independent seedlings, with about 400-600 cells examined per marker combination for endosome counting.

To understand the role of PICALMs in ERECTA internalization, we assessed the formation of ERECTA-YFP-positive BFA bodies in ’wild-type’ (*ERECTApro::ERECTA-YFP* in *er-105*) seedlings under BFA treatment (Fig. 5A), compared with the *picalm1a;1b;4a;4b* quadruple mutant background. In WT epidermal cells, ERECTA-YFP formed numerous BFA bodies, particularly in stomatal lineage ground cells (SLGCs) and developing pavement cells (PCs), suggesting elevated endocytic activity in these cell types (Fig. 5A upper panel; 5B). Strikingly, in the *picalm1a;1b;4a;4b* mutant background, the number of ERECTA-YFP-positive BFA bodies is significantly reduced (Fig. 5A lower panel; 5C), suggesting a marked impairment in ERECTA internalization. Neither transcript-nor protein levels of ERECTA-YFP, as well as the endogenous *ERECTA* levels, are affected by the *picalm1a;1b;4a;4b* mutations (fig. S4A-C). Thus, the observed ERECTA internalization defects in higher-order *picalm* mutants are not due to altered ERECTA transcription or protein levels. To further address the mechanistic basis of these internalization defects, we performed a pharmacological assay using the CME inhibitor ES9-17 (*65*). In wild type, ES9-17 significantly reduced the numbers of ERECTA-YFP positive BFA bodies, confirming the effective inhibition of CME (fig. S5). In contrast, the *picalm1a;1b;4a;4b* mutant showed no further reduction upon ES9-17 treatment (fig. S5), indicating that ERECTA internalization is already compromised and insensitive to CME inhibition. These results demonstrate that PICALMs are required for CME-dependent entry steps of ERECTA trafficking.

**Fig. 5.**
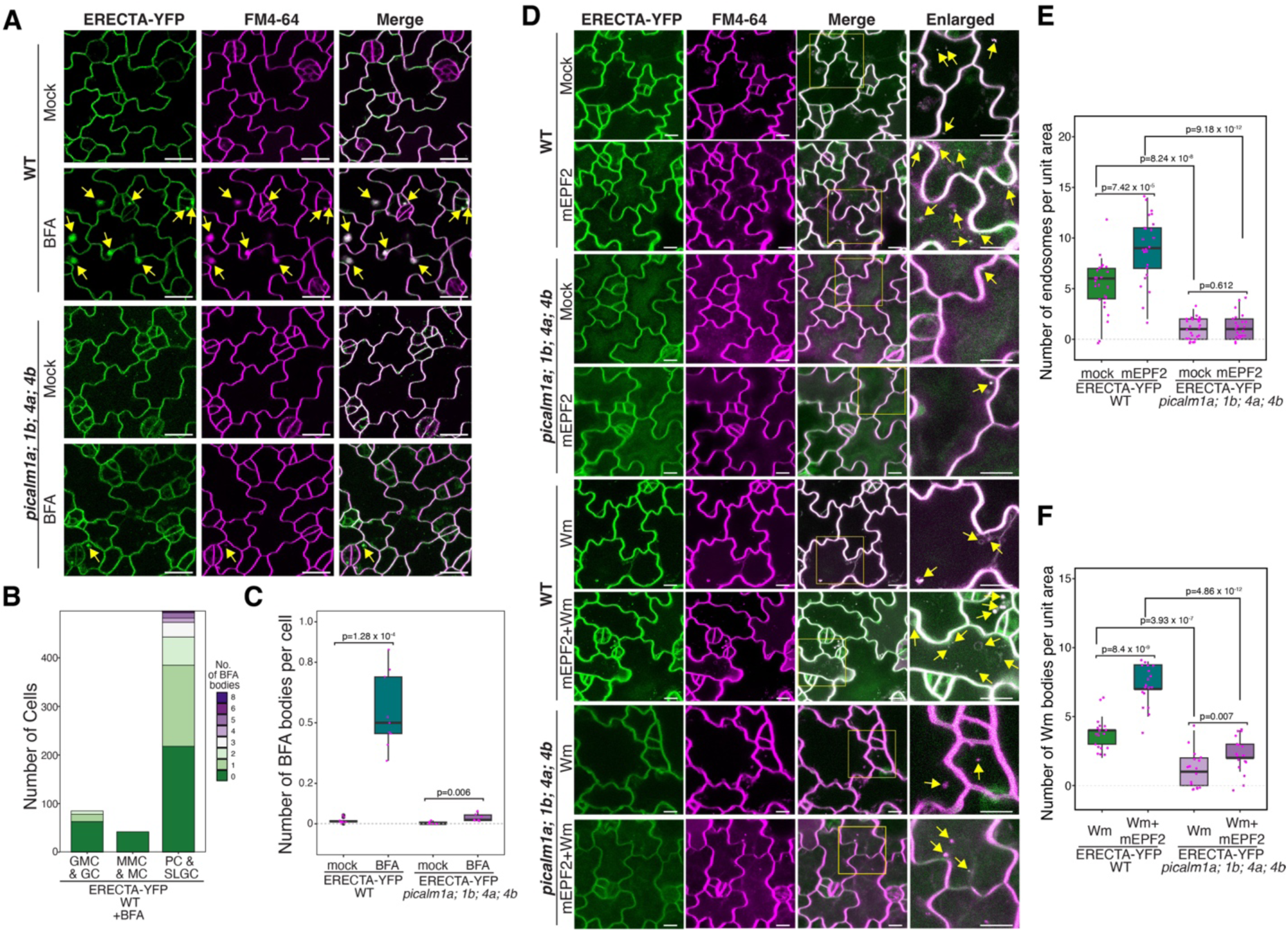
PICALM proteins regulate ERECTA internalization. **(A)** Confocal microscopy images of ERECTA-YFP in WT and *picalm1a;1b;4a;4b* mutant in the abaxial cotyledon epidermis at 4 days after germination under mock or BFA treatment. ERECTA-YFP in green and FM4-64 membrane dye in magenta. Yellow arrows, ERECTA-YFP–positive BFA bodies. **(B)** Quantification of the number of cells with BFA bodies across stomatal lineage stages in WT. ERECTA-YFP endocytic activity is enriched in pavement cells (PCs) and stomatal lineage ground cells (SLGCs). n = 9 independent seedlings, with 858 cells examined for BFA body counting. **(C)** Quantitative analysis of the number of BFA bodies per cell under mock and BFA treatments in WT and *picalm1a;1b;4a;4b* mutant. Box plot shows the interquartile range (IQR; 25th-75th percentile), horizontal line indicates the median, whiskers extend to 1.5× IQR, and individual data points are overlaid. Welch’s two-sample t-test was performed. n = 9 independent seedlings, with about 800-1000 cells examined per treatment for BFA body counting. **(D)** Confocal microscopy images showing ERECTA-YFP internalization in response to mature EPF2 (mEPF2) peptide, Wortmannin (Wm), or their combination in WT and *picalm1a;1b;4a;4b* mutant. Yellow arrows, endocytic vesicles or Wm bodies. **(E)** Quantitative analysis of mEPF2-induced ERECTA-YFP puncta in WT and *picalm1a;1b;4a;4b* mutant shows impaired ligand-triggered ERECTA endocytosis in the mutant. Box plots show the number of intracellular endosomes per unit area, with Welch’s two-sample t-test performed for pairwise comparisons (n = 25 independent seedlings). **(F)** Quantification of ERECTA-YFP positive Wm bodies per cell in WT and *picalm1a;1b;4a;4b* mutant. Wm-induced accumulation is significantly reduced in the mutant, indicating defective trafficking to late endosomes. Box plots show the number of Wm bodies per unit area, with Welch’s two-sample t-test performed for pairwise comparisons (n = 18 independent seedlings).

We subsequently tested whether PICALMs are required for the ligand-induced internalization of ERECTA. In WT, treatment with exogenous mature EPF2 (mEPF2) peptide triggered cytoplasmic accumulation of ERECTA-YFP puncta, overlapping with FM4-64, suggesting a ligand-activated endocytosis (Fig. 5D, 5E). This response was greatly reduced in *picalm* mutants, where ERECTA-YFP remained predominantly plasma membrane-localized (Fig. 5D, upper panel; 5E). Wm treatment induced ERECTA-YFP accumulation in enlarged late endosomes (Wm bodies) in WT, and co-treatment with mEPF2 and Wm further enhanced this effect. In contrast, the *picalm* quadruple mutants showed minimal ERECTA-YFP accumulation in Wm bodies under both treatment conditions (Fig. 5D, lower panel; 5F), indicating impaired endocytic trafficking to late compartments. Together, these findings show that PICALMs are required for both constitutive and ligand-induced endocytosis of ERECTA, underscoring their role as endocytic adaptors essential for receptor internalization and intracellular trafficking.

Lastly, to assess whether internalized ERECTA proteins are destined to the lytic vacuole, we performed a pharmacological assay by treating seedlings with Concanamycin A, an inhibitor of vacuolar H^+^-ATPase that raises vacuolar pH and preserves ERECTA-YFP signals (*66*). Concanamycin A treatment led to clear accumulation of ERECTA-YFP inside the lytic vacuole (fig. S6A, B, tonoplast marked by FM4-64), consistent with vacuolar trafficking previously observed for ERL1 (*32*). In contrast, vacuolar ERECTA-YFP was largely absent in the *picalm1a;1b;4a;4b* mutant (fig. S6A, B), further highlighting the critical role of PICALMs for ERECTA internalization.

### *PICALMs* redundantly enforce stomatal patterning by maintaining ERECTA homeostasis

The association and colocalization of PICALM family members with ERECTA during CME (Figs. 1 to 4), together with impaired ERECTA internalization observed in *picalm* mutant (Fig. 5), prompted us to investigate their functional roles in ERECTA-mediated stomatal development. For this purpose, we first examined if the *picalm* mutants exhibit stomatal patterning defects. None of the single loss-of-function mutants of PICALMs that we tested (*picalm1a, picalm1b, picalm2a, picalm2b, picalm4a, picalm4b, picalm5a*, and *picalm5b*) exhibited any discernible phenotypes in stomatal development (fig. S8), indicating that these *PICALMs* are functionally redundant. Notably, *picalm3* mutant showed a weak stomatal patterning defect (Fig. 6A). Consistent with the synergistic nature of their interactions, *picalm* mutants displayed escalating stomatal patterning defects, characterized by increased stomatal number and clustering, with the higher-order mutants (*picalm1a;1b;4a;4b*, *picalm1a;1b;3;4a;4b*) exhibiting a more severe phenotype than the single (*picalm3*), double (*picalm1a;1b*, *picalm4a;4b*), or triple (*picalm3;4a;4b*) mutants (Fig. 6A). Importantly, the stomatal clustering phenotype resembles that of the *erecta*-family mutants (*12*) (fig. S9). Quantitative analysis confirmed a significant increase in stomatal index and clustering across multiple *picalm* mutants (Fig. 6B, 6C), indicating that *PICALMs* are required for proper stomatal patterning.

**Fig. 6.**
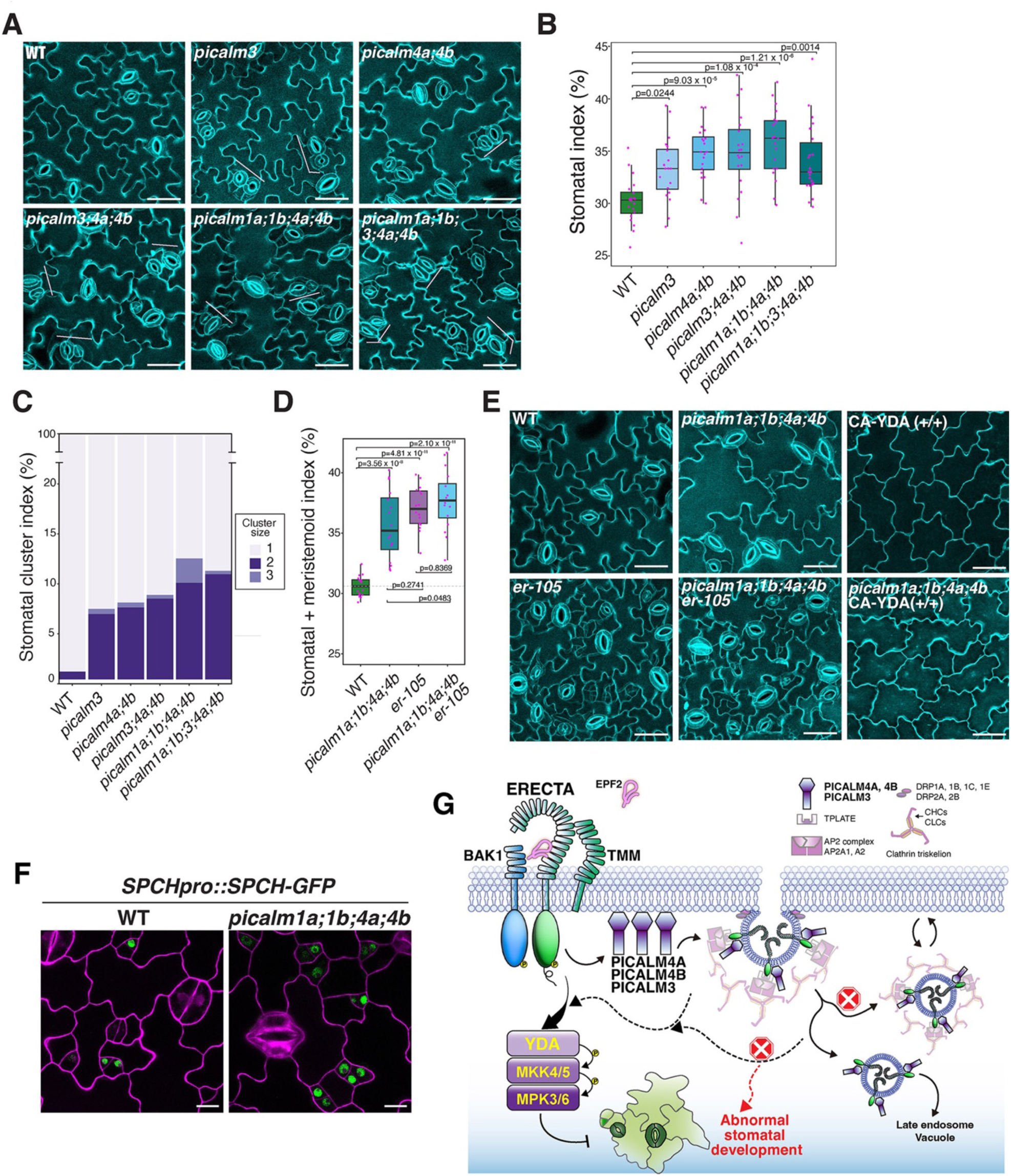
*PICALMs* are required for proper stomatal patterning by ERECTA signaling pathway. **(A)** Confocal images of abaxial epidermis from wild-type (WT) and selected *picalm* mutants. WT plants exhibit normal stomatal development, whereas *picalm3, picalm4a;4b, picalm3;4a;4b, picalm1a;1b;4a;4b,* and *picalm1a;1b;3;4a;4b* mutants exhibit progressively increased stomatal index with varying degrees of stomatal clustering. Scale bars: 40 μm. **(B)** Quantification of stomatal index in WT and selected *picalm* mutants. One-way ANOVA was performed followed by Tukey’s HSD test. n = 20 cotyledons from independent seedlings. **(C)** Stomatal cluster index analysis in WT and *picalm* mutants. Stomatal clusters were categorized as: 1 = one normal stoma, 2 = paired stomata, 3 = cluster of three or more stomata. n = 20 independent cotyledons from independent seedlings. **(D)** Quantification of combined stomatal and meristemoid index in WT and selected *picalm* mutants. The observed non-additive genetic interaction between *er-105* and *picalms* is consistent with PICALMs acting in the same genetic pathway as ERECTA. n = 16 independent cotyledons from distinct seedlings. **(E)** Confocal microscopy images of abaxial cotyledon epidermis from WT, *er-105*, *picalm1a;1b;4a;4b*, *picalm1a;1b;4a;4b;er-105*, *picalm1a;1b;4a;4b;CA-YDA (+/+)*, and the *CA-YDA(+/+)* line. Constitutively active YDA suppresses stomatal formation in both WT and *picalm1a;1b;4a;4b*. Scale bars: 40 μm. **(F)** SPCH-GFP accumulation in WT and *picalm1a;1b;4a;4b*. More cells exhibit elevated SPCH-GFP signals in *picalm* quadruple mutants. Scale bars: 10 μm. **(G)** Proposed model of PICALM-mediated ERECTA trafficking in stomatal development. Upon EPF2 perception, ERECTA is internalized through CME involving adaptor complexes containing PICALM3, PICALM4A, and PICALM4B, along with the AP2 and TPLATE complexes. Internalized receptors signal through the YDA-MKK4/5-MPK3/6 cascade to restrict SPCH activity and ensure proper stomatal development. In *picalm* mutants, defective CME impairs ERECTA internalization, resulting in disrupted stomatal patterning.

To explore the genetic relationship between PICALMs and ERECTA, we generated a *picalm1a;1b;4a;4b;er-105* quintuple mutant and compared its phenotype to the *erecta* null mutant, *er-105*. Both mutants exhibited comparable increases in stomatal and meristemoid indices, with no significant additive effects in the quintuple mutant (Fig. 6D, 6E). To further address whether PICALMs function within the ERECTA signaling pathway, we investigated whether the activation of the downstream MAPK cascade could mask the stomatal clustering phenotype of the higher-order *picalm* mutant. Indeed, constitutively active YDA (CA-YDA) conferred the ‘pavement-cell-only’ epidermis in *picalm1a;1b;4a;4b* seedlings, just like in wild type (Fig. 6E). Moreover, CA-YDA partially rescued the dwarf phenotype of the higher-order *picalm* mutant plants (fig. S12). These results suggest that PICALMs act upstream of the MAPK cascade within the ERECTA signaling pathway. It is known that the ERECTA-MAPK signaling pathway directly targets SPEECHLESS (SPCH) protein stability (*17, 67*). We thus examined whether the higher-order *picalm* loss-of-function mutations confer the dysregulation of SPCH-GFP protein levels. As shown in Fig. 6F, more stomatal precursor cells exhibited high SPCH-GFP signals in *picalm1a;1b;4a;4b* seedling epidermis, suggesting that impaired PICALM function dampens the canonical ERECTA-YDA-MAPK output.

To conceptualize this mechanism, we propose a model in which PICALMs, particularly PICALM4A, 4B, and PICALM3, function as cargo-selective adaptors that recruit ERECTA into CME together with the AP2 and TPLATE complexes (Fig. 6G). Upon EPF2 perception, ERECTA internalization via this PICALM-dependent route enables robust activation of YDA-MKK4/5-MPK3/6 cascade to restrict SPCH activity and maintain proper stomatal patterning (Fig. 6G). In *picalm* mutants, defective CME impairs ERECTA internalization, resulting in abnormal stomatal development.

## Discussion

In this study, we comprehensively mapped the ERECTA signaling network in *Arabidopsis* using a dual proteomic strategy, ET-AP and TbID-PL. Our findings not only recovered stable ERECTA receptor complex components but also identified a dynamic cohort of transient, low-abundance interactors, notably CME components such as PICALMs. Functional characterization of PICALMs demonstrates their essential role in ERECTA internalization and protein homeostasis, with higher-order *picalm* mutants phenocopying the excessive stomatal number and clustering observed in *ERECTA*-family loss-of-function mutants. A previous study has demonstrated ligand-induced endocytosis of the ERECTA-family member, ERL1(*32*); however, the specific adaptors facilitating this process remained unknown. Our work identifies the components of CME that recognize ERECTA as a cargo and further emphasizes that CME-mediated receptor dynamics are indispensable for ERECTA receptor kinase-mediated stomatal patterning through the YDA MAPK cascade, which eventually targets SPCH (Fig. 6).

While proteomic profiling has been increasingly utilized to uncover receptor kinase interactors (*68, 69*), many challenges still persist that hamper the identification of low-abundance or transient interactors without disrupting the native protein complex in the membrane environment. Our optimized, combinatorial approach using ET-AP and TbID-PL enabled the detection of both known components of the ERECTA signaling pathways as well as previously undetected components. Technically, the exclusive detection of TMM by ET-AP, but not by TbID-PL (Fig. 1), highlights the preservation of the native plasma membrane environment during proximity labeling, ensuring the reliability of our *in vivo* proteomic data. Our refined TbID-PL enriched low-abundance signaling components like BSK1 and YDA (Fig. 2), which are otherwise difficult to detect using conventional methods. ERECTA regulates diverse roles in development and environmental response (*9, 70*), and as such, the identification of additional interactors not previously associated with the ERECTA pathways, such as LIK1 and TMK4 (Figs. 1 and 2), may pave the way for future investigations of signal crosstalk.

The exclusive identification of CME components by TbID-PL highlights the transient nature of their interactions with the receptor kinase (Fig. 1). For example, the PICALM members, PICALM1A, PICALM3, and PICALM4A/4B, associate with ERECTA to varying degrees and exhibit unequal redundancy in enforcing stomatal patterning (Fig. 6, A to C). Unlike their metazoan counterparts, which comprise only four members in humans, the PICALM family is vastly expanded in plants, with 18 members in *Arabidopsis* (*71*). These AtPICALMs contain a conserved ANTH (AP180 N-terminal homology) domain, which is essential for binding to phosphatidylinositol lipids and clathrin, facilitating cargo endocytosis (*71*). However, their slight structural differences may enable them to interact with the same cargo with different affinities, as we observed for ERECTA (Fig. 3D), and/or alternatively, with distinct sets of membrane cargos or adaptor proteins. In fact, a recent study suggests that PICALM1A and PICALM1B interact with R-SNAREs and clathrin at the plasma membrane, where they regulate R-SNARE recycling to support plant growth (*60*). Additionally, PICALM5A and PICALM5B are required for CME during pollen tube elongation by maintaining tip localization of ANXUR, a receptor kinase sensing cell wall integrity (*72*). PICALM4A (also known as ECA4) has also been shown to act synergistically with another clathrin adaptor protein EPSIN to support apical cell growth in pollen tubes (*73*). Moreover, PICALM4A/ECA4 has been implicated in the endocytosis of both the brassinosteroid receptor BRI1 and the ABA exporter ABCG25 in roots (*74*). In addition to stomatal patterning defects, we observed that higher-order *picalm* mutations confer broader developmental defects, including reduced fertility and impaired root elongation (fig. S10C and fig. S11). Therefore, these PICALM subfamily members identified through TbID-PL likely regulate additional clients beyond ERECTA.

We observed cell-type specific differences in ERECTA internalization across the epidermis, with notably high internalization activity in developing stomatal lineage ground cells (SLGCs) and pavement cells, and low levels in meristemoids (including MMCs) and GMCs (Fig. 5A and 5B). It is well established that ERECTA perceives EPF2 and restricts the initiation of stomatal cell lineages via MAP kinase-mediated degradation of SPCH proteins (*16, 17, 19, 67, 75*). High activity of ERECTA internalization in the SLGCs and pavement cells suggests that ERECTA signaling is actively restraining their division potential, thereby limiting their re-entry into stomatal cell lineages. This view is supported by the recent report that SPCH protein levels determine whether SLGCs undergo asymmetric division or differentiate into mature pavement cells, a decision impacted by EPF-signaling in neighboring cells (*76*). The EPF-ERECTA-family ligand-receptor system is known to function in both autocrine and juxtacrine signaling (*77, 78*). How PICALM-mediated endocytosis influences these distinct modes of peptide-receptor signaling is an interesting future question. We also observed that while ERECTA colocalized with PICALM4A at the plasma membrane in stomatal lineage cells, it was notably absent from the cell plates of dividing GMCs (Fig. 3C, Late GMC). In contrast, another ERECTA-family receptor, ERL1, was present at the cell plates and colocalized with PICALM4A in these dividing cells (fig. S7A). This spatial divergence suggests that PICALM4A may differentially associate with ERECTA-family members depending on subcellular context or developmental stage. Consistent with this overlap, our Co-IP experiments show that ERL1 also interacts with PICALM4A *in vivo* (fig. S7B). Together with the enhanced stomatal clustering phenotype observed in the *picalm1a;1b;4a;4b;er-105* quintuple mutant compared to *er-105* alone, these findings further support the notion that PICALMs may mediate trafficking of multiple ERECTA-family receptor kinases.

During the EPF ligand recognition by ERECTA family receptors, the EPF1/2 antagonist Stomagen/EPFL9 triggers stalling of ERL1 in the endoplasmic reticulum, coupled with reduced endocytosis (*32*). The process likely involves the endoplasmic reticulum-resident chaperone complex SDF2-ERdj3B-BiP, which facilitates proper translocation of ERECTA-family RKs from the endoplasmic reticulum to plasma membrane (*79, 80*). By analogy, defects in ERECTA internalization, such as observed in *picalm* mutants, could similarly impede receptor turnover at the plasma membrane, thereby creating a “traffic jam” that interferes with normal receptor homeostasis and proper signaling output.

Our observations also imply that the subcellular dynamics of ERECTA may be coupled with its post-translational modifications. Specifically, EPF2-induced activation of ERECTA triggers its phosphorylation by coreceptor BAK1, followed by ubiquitination via the U-box E3 ligases PUB30 and PUB31, to attenuate signaling (*24*). This process likely promotes ubiquitination-dependent internalization of ERECTA, consistent with prior findings that ubiquitination promotes BRI1 endocytosis and enables its recognition at the trans-Golgi network/early endosomes (TGN/EE) for vacuolar targeting (*81*). Consistently, a very recent study identified specific ubiquitination sites of ERECTA and further demonstrated that ligand-activated ubiquitination triggers the endocytosis of ERECTA (*82*). In that study, high accumulation of the ubiquitination-deficient mutant version of ERECTA at the plasma membrane caused inhibition of stomatal development, indicative of the overly activated ERECTA signaling due to failed attenuation. In contrast, we here revealed that the loss-of-function mutations of *PICALM* subfamily members perturb the endocytosis of ERECTA and result in stomatal clustering (Figs. 5 and 6), indicating that the ERECTA signaling pathway is compromised in the absence of *PICALMs*. This apparent paradox, whereby defective endocytosis of ERECTA leads to opposite signaling outcomes, raises interesting questions about the multiple layers of regulation imposed by receptor trafficking. We speculate that the activation of the ERECTA receptor complex requires the formation of membrane subdomains reinforced by the PICALM-mediated cargo sorting and vesicle formation. In such a case, PICALMs may facilitate the assembly of signaling-competent receptor-coreceptor complexes within specific plasma membrane nanodomains. Alternatively, this early activation process can be mechanistically distinct from the later steps of ubiquitination-mediated internalization and eventual degradation of ERECTA that attenuates signaling. It is likely that different endocytic routes downstream of PICALMs other than ubiquitination direct ERECTA toward distinct subcellular fates. Additionally, PICALMs may generally facilitate endocytic trafficking, and if so, the *picalm* higher-order mutants could disrupt additional factors that impact ERECTA signaling as well as stomatal development.

These findings collectively support the model that ERECTA signaling output is not solely dictated by receptor abundance, but rather by its dynamic partitioning into specialized membrane domains and intracellular compartments. Further understanding of ERECTA receptor trafficking will provide a foundation for future research into how membrane dynamics shape cellular differentiation and patterning. Looking forward, our proteomic atlas offers a framework for dissecting other RLK networks, particularly those governing environmental responses where trafficking may modulate signaling plasticity.

## Limitations of the study

Due to technical limitations, our TbID-PL approaches did not include ligand-induced proteomic profiling with exogenously synthesized EPFs, making it challenging to distinguish receptor dynamics before and after ligand perception. However, under the native conditions we used, plant cells endogenously secrete EPF peptides, leading to continuous inhibition of excessive stomatal development. Therefore, our profiling most likely captured a pool of ERECTA-family receptors, including both ligand-activated and inactivated states. Indeed, ligand-induced interactors such as BAK1 were also identified, supporting the validity of our approach.

## Materials and Methods

### Plant materials and growth conditions

*Arabidopsis* Col-0 was used as the wild-type control in this study. The following materials have been published previously: *er-105*, *erl1-2, erl2-1, er-105;erl1-2, er-105;erl2-1, erl1-2;erl2-1*, *er-105;erl1-2;erl2-1* (*11*); *ARA6pro::ARA6-mRFP, ARA7pro::mRFP-ARA7, SYP22pro::SYP22-mRFP, SYP43pro::SYP43-mRFP* (*83*), CA-YDA (*84*), *SPCHpro::SPCH-GFP* (*85*). The following Arabidopsis mutants were obtained from Arabidopsis Biological Resource Center (ABRC): *picalm1a* (SALK_043625), *picalm1b* (GABI_026G05), *picalm2a* (WiscDsLoxHs212_04A), *picalm2b* (SALK_032696), *picalm3* (WiscDsLoxHs184_04D), *picalm4a* (WiscDsLox442D1), *picalm4b* (SAIL_418_A409), *picalm5a* (SALK_080752), *picalm5b* (SALK_127035). Higher-order mutants of *PICALM* family were generated through genetic crosses, including double mutant *picalm1a;1b*, *picalm4a;4b*, triple mutant *picalm3;4a;4b*, quadruple mutant *picalm1a;1b;4a;4b*, and quintuple *picalm1a;1b;3;4a;4b*. Arabidopsis seeds were surface sterilized with 2.5 % sodium hypochlorite solution, stratified at 4 °C in darkness for two days, and then germinated on half-strength Murashige-Skoog (1/2 MS) medium plates for 8-10 days before imaging or transplanting to soil pots. The 1/2MS medium (pH 5.7) contains 1.0 % sucrose, 1x Gamborg′s Vitamins (Sigma, Cat # G1019), and 0.75 % agar. Plants were grown in a controlled growth room under a long-day cycle of 16 hr light (22 °C) and 8 hr dark (20 °C).

### Plasmid constructions and plant transformation

The *ERECTA-TurboID* construct pPFB11 (*ERECTApro::ERECTA-HA-TbID*) and the negative control pPFB36 (*ERECTApro::Lti6B-HA-TbID*) were generated using a MultiSite Gateway cloning strategy. Entry vectors containing the *ERECTA* promoter and entry vectors containing either *ERECTA-HA-TbID* or *Lti6B-HA-TbID* fusions were recombined into the R4pGWB501 destination vector (*86*). For protoplast expression, genomic coding sequences of *PICALM3*, *PICALM4A*, and *PICALM4B* were PCR-amplified from Col-0 genomic DNA and cloned into the pYBA-1132 vector (35S::C-terminal EGFP, NCBI Accession: KF876796) via *BamHI* and *KpnI* restriction sites. Additionally, the genomic coding sequence of *ERECTA* was cloned into the pHBT-2xHA vector using *BamHI* and *StuI* restriction sites. To generate constructs for the AlphaScreen assay, coding sequences (CDSs) of *PICALM* family genes (*PICALM1A*, *PICALM2B*, *PICALM3*, *PICALM4A*, *PICALM4B*, *PICALM5A*, *PICALM6*, *PICALM8*, *PICALM9B*, *PICALM10B*) and the cytoplasmic kinase domain of *ERECTA* (ER-KD) were PCR-amplified from Col-0 cDNA. The amplified CDSs were cloned into pENTR/D-TOPO entry vectors and recombined into the destination vectors pEU-E01-GW-bls-STOP (*87*) and pEU-E01-AGIA-GW-STOP (*88*). GFP expressed from pEU-E01-AGIA-GW-STOP served as a negative control. Two additional constructs used for subcellular localization and complementation experiments, *PICALM3pro::PICALM3-TagRFP-pGWB559* and *PICALM4apro::PICALM4a-mRFP-pGWB1*, were generated by Gateway cloning. A full list of plasmids and primers used is provided in Tables S1 and S2. Stable transgenic plants were generated using *Agrobacterium tumefaciens* (GV3101::pMP90) -mediated floral dipping method (*89*). At least 20 independent T1 transformants were analyzed for each construct. Stable transgenic lines were identified by segregation analysis for single-locus insertion (3:1 segregation ratio).

### Confocal microscopy imaging

Confocal imaging for stomatal phenotyping was conducted using a Leica SP5-WLL microscope equipped with a 20x dry objective lens. Cotyledons from 8 days post-germination (dpg) *Arabidopsis* seedlings were stained with 0.1 mg/mL propidium iodide (PI) (Sigma, Cat. # P4864) for 1 min to visualize cell outlines. The abaxial epidermis was imaged using an excitation wavelength of 561 nm and an emission detection range of 590-659 nm. Z-stack images spanning the full depth of the epidermal cell layers were captured and processed using Fiji/ImageJ software (http://fiji.sc/Fiji). Maximum intensity projections were generated, and representative images were false-colored and adjusted for brightness and contrast using Adobe Photoshop 2022. For subcellular colocalization and internalization analyses, fluorescent signals were captured using the Leica SP5-WLL with a 63x water immersion objective lens, or the Leica Stellaris 8 FALCON microscope with 40x or 63x oil immersion objective lens operated in Tau Gating mode. Fluorescence excitation/emission settings were optimized for specific fluorophores as follows: YFP (514 nm excitation, 520-550 nm emission), mRFP (584 nm excitation, 600–650 nm emission), and TagRFP (555 nm excitation, 565–650 nm emission).

### Pharmacological treatment

For exogenous chemical treatments, 4 dpg Arabidopsis seedlings were pre-stained in ultrapure water containing 5 µM FM4-64 (Thermo Scientific, Cat. # T3166) for 10 min. Following staining, seedlings were immersed in ½ MS liquid medium supplemented with 30 µM BFA (Sigma, Cat. # B7651, 30 mM stock in ethanol), 30 µM Wortmannin (Sigma, Cat. # W1628, 30 mM stock in DMSO), or 50 µM cycloheximide (Sigma, Cat. # C4859, 100 mg/mL stock in DMSO). Seedlings were vacuum infiltrated for 1 min to facilitate chemical uptake and subsequently incubated for 60 min prior to imaging. For EPF2 ligand-induced internalization analyses, recombinant mEPF2 peptide was purified and refolded to retain its bioactivity (*16*). After confirming its bioactivity, the peptide was diluted to 5µM using 1/2MS liquid medium (*49*), with or without 30 µM Wortmannin. Seedlings were incubated for 90 min followed by 5 µM FM4-64 staining before imaging. For ES9-17 (Sigma, Cat. # SML2712) treatment, cotyledons of 4-day-old seedlings were first stained with 5 µM FM4-64 for 30 min, then transferred to either a mock solution (½ MS liquid medium supplemented with 0.1% DMSO, 50 µM cycloheximide, and 50 µM BFA) or to an ES9-17 solution (½ MS liquid medium supplemented with 100 µM ES9-17, 50 µM CHX, and 50 µM BFA). Seedlings were vacuum-infiltrated for 1 min and subsequently incubated for 1 h in the mock or ES9-17 solution before imaging. For Concanamycin A (Adipogen Life Sciences, Cat. # BVT-0237-C100) treatment, cotyledons of 4-day-old seedlings were stained with 5 µM FM4-64 for 30 min, then rinsed and incubated in ultrapure-water for 6 h to washout the excess dye. Seedlings were then vacuum-infiltrated with 1 µM Concanamycin A in ½ MS liquid medium, followed by an additional 6 h incubation before imaging.

### Sample preparation for proximity labeling

TurboID-based proximity labeling assays were conducted following previously described procedures with modifications (*40, 41, 43*). Seeds of *ERECTApro::ERECTA-HA-TbID*, *ERECTApro::Lti6B-HA-TbID* transgenic lines, together with *er-105* mutant were surface-sterilized with 70% ethanol and sown on filter paper (90 mm, Cytiva Whatman™, Cat # 09-853A) placed on 1/2 MS medium plates. Five-day-old seedlings grown under long-day conditions were harvested and transferred to a flask for biotin treatment. To optimize labeling conditions, a range of biotin solutions (0, 5, 20, 50, 100, and 250 µM) and incubation durations (0, 15 min, 30 min, 1 hr, 2 hr, 3 hr) were tested. Optimal conditions for the ERECTA-TurboID PL assay were determined to be an incubation with 50 µM biotin for 1 hr at room temperature. To terminate the PL reaction, seedlings were rinsed five times with ice-cold water to remove excess biotin, gently blotted dry with laboratory Kimwipes, and immediately frozen in liquid nitrogen. Frozen samples were ground into a fine powder under liquid nitrogen for protein extraction. All experiments were performed in three independent biological replicates.

Approximately 2.0 mL of the finely ground samples were homogenized in 2.5 mL of ice-cold protein lysis buffer (100 mM Tris pH 7.5, 150 mM NaCl, 10 % glycerol, 20 mM Sodium fluoride, 1.5 mM Sodium orthovanadate, 2 mM Sodium molybdate, 1 mM PMSF, 1 % Triton X-100, 1x Protease inhibitor cocktail) in a 5.0 mL LoBind tube (Eppendorf, 0030108302). The mixture was incubated on a rotor wheel at 4 °C for 30 min and then sonicated four times at 4 °C for 30s with 90s intervals on a high setting using a Bioruptor 300 (Diagenode). Next, the homogenized samples were centrifuged at 13,000 rpm for 10 min at 4 °C to remove cell debris. Approximately 2.5 mL of the supernatant was processed through a PD-10 desalting column (Cytiva, Cat # 17-0851-01) using the gravity flow protocol to remove excess free biotin. Approximately 3.5 mL of protein extract was collected, and the protein concentration was determined using the Bradford assay (Bio-Rad). The protein extracts were divided equally for the analysis of the ERECTA proximitome and interactome. For proximitome, 1.7 mL of lysate (approximately 5.5 mg of protein) was transferred to a new 2.0 mL LoBind tube (Eppendorf, 0030108450) and incubated with 100 μL of pre-equilibrated streptavidin magnetic beads (Thermo Scientific, Cat # 88817) at 4°C for 18 hr on a rotor wheel. The streptavidin magnetic beads, enriched with biotinylated proteins, were sequentially washed three times with protein lysis buffer, followed by washes with 1 M KCl, 0.1 M Na_2_CO_3_, 2 M urea, and 50 mM Tris-HCl (pH 8.0). One-tenth of the beads were used for Western blot analysis, and the remaining beads were frozen for subsequent analysis. Similarly, the second portion (5.5 mg of protein), was incubated with 100 μL of pre-equilibrated anti-HA magnetic beads (Thermo Scientific, Pierce™, Cat # 88837) under the same conditions. The anti-HA magnetic beads were washed three times with protein lysis buffer and three times with 50 mM Tris-HCl (pH 8.0). One-tenth of the beads were used for immunoblot analysis, and the remaining beads were frozen for subsequent analysis. Immunoblot analysis showed that ERECTApro::Lti6B-HA-TbID accumulated at very low levels, resulting in minimal α-HA and streptavidin signals. Screening of 18 independent lines confirmed consistently weak expression, likely due to limitations within the Lti6B-HA-TbID construct.

### LC-MS sample preparation

Biotinylated proteins bound to streptavidin magnetic beads were processed for mass spectrometry analysis using an on-bead trypsin digestion protocol. The streptavidin magnetic bead samples, previously stored at -80 °C, were thawed and washed three times with 50 mM Tris-HCl buffer (pH 8.0) to remove excess detergents and salts. An equal volume of trifluoroethanol (TFE) was added to denature the proteins and expose trypsin cleavage sites. Tris(2-carboxyethyl) phosphine (TCEP) was added to a final concentration of 5 µM, and the mixture was incubated at 55°C for 45 min to reduce disulfide bonds. After cooling to room temperature, iodoacetamide (15 mM final concentration) was added to alkylate cysteine residues and prevent the reformation of disulfide bonds. This reaction was conducted in the dark at room temperature for 30 min and quenched with 7 mM dithiothreitol (DTT). For trypsin digestion, the mixture was diluted with 50 mM Tris-HCl (pH 8.0) containing 2 mM CaCl_2_ and 2.0 µg MS-grade trypsin (Thermo Scientific, Cat # 90058). The digestion reaction was incubated at 37 °C for 5 hr with end-to-end rotation and terminated by adding formic acid to a final concentration of 1% (v/v). Protein complexes enriched with anti-HA magnetic beads were eluted by washing the beads three times with 0.1 M glycine (pH 2.5), followed by separation of the supernatant from the bead pellet. The eluate was neutralized with 1M Tris-HCl buffer (pH 8.0) and subjected to the same trypsin digestion protocol as described above. After trypsin digestion, the supernatant was separated from the bead pellet and concentrated using a vacuum concentrator (Vacufuge plus, Eppendorf). The sample was desalted using C18 pipette tips (Thermo, Cat # 60109-412), dried in a SpeedVac, and resuspended in buffer (5 % acetonitrile, 95 % LC-MS grade water with 0.1 % formic acid) for liquid chromatography-mass spectrometry (LC-MS) analysis.

### Mass spectrometry data analysis

TbID-PL and ET-AP mass-spectrometry experiments were performed with three biological replicates. For TbID-PL, each biological replicate was analyzed with two LC-MS injections, yielding a total of 18 raw files. For ET-AP, each biological replicate was analyzed with three LC-MS injections, yielding a total of 27 raw files. Raw MS/MS spectra were acquired using a Thermo Orbitrap Fusion Lumos mass spectrometer and processed using Proteome Discoverer (version 2.3) following the PWF_Basic_SequestHT workflow with the Percolator node for peptide spectrum match (PSM) validation. The processing settings included trypsin digestion with up to two missed cleavages, static modifications of carbamidomethyl cysteine, dynamic modifications of oxidized methionine, and protein N-terminal modifications of acetylation and/or methionine-loss. Mass spectra were searched against the Arabidopsis thaliana (ARATH) reference proteome, TurboID, and common contaminants (downloaded from https://www.uniprot.org). High-confidence PSMs, peptides, and proteins were all filtered at a false discovery rate of < 5%. Individual RAW files were processed to generate .msf files, which were reprocessed together using the CWF_Basic workflow with Merge Mode set to “Do Not Merge” to generate a single consensus file for comparative analysis. Proteins significantly associated with ERECTA bait were identified by calculating log_2_ fold-changes and Z-scores based on observed PSMs between experimental and control conditions, using the R package diffprot (https://github.com/rachaelcox/diffprot.git).

### Protein Interaction network construction

Proteins significantly enriched in the TurboID-PL and ET-AP datasets (Data S1, S2, and S3) were selected and mapped to the STRING database (https://string-db.org/) using a medium confidence score threshold (interaction score ≥ 0.4). The resulting protein-protein interaction (PPI) network was visualized in Cytoscape (version 3.9.1) (*90*), with edges representing both experimentally determined and computationally predicted interactions (e.g., gene neighborhood, fusions, and co-occurrence). Node colors were used to distinguish the proteomic approaches, while node size reflected interaction degree. Functional clusters were manually curated to highlight key biological processes, including internalization, exocytosis, and stomatal development, based on Gene Ontology (GO) annotations and pathway analysis.

### Protoplast transfection assay

The isolation and transfection of Arabidopsis protoplasts were performed with minor modifications to the protocol as previously described (*91*). Arabidopsis Col-0 plants were grown under short-day conditions (8 hr light at 22 °C/ 16 hr dark at 20 °C). For protoplast isolation, rosette leaves from 4-week-old plants were cut into 0.5-1.0 mm strips and digested in an enzymatic solution containing 1 % Cellulase R10 (Yakult Honsha, Japan), 0.2 % Macerozyme R10 (Yakult Honsha, Japan), 0.4 M mannitol, 20 mM KCl, 20 mM MES, 10 mM CaCl_2_, 5 mM β-mercaptoethanol, and 0.1 % BSA. Digestion was carried out with gentle shaking (40 rpm) for 3 hr in the dark. The protoplasts were collected by filtering the solution through a 70 µm nylon mesh (Corning, Cat # 352350) and washed three times with W5 solution (154 mM NaCl, 125 mM CaCl_2_, 5 mM KCl, 5 mM glucose, 2 mM MES, pH 5.7) to remove residual enzymes. The protoplasts were resuspended in MMG solution (400 mM mannitol, 15 mM MgCl_2_, 4 mM MES, pH 5.7) for transfection. For PEG-mediated transfection, 200 μL of protoplast suspension (5 × 10^5^ cells/mL) was mixed with 10-15 µg of plasmid DNA in a 1.5 mL Eppendorf LoBind tube. The mixture was incubated with an equal volume of freshly prepared PEG solution (40% PEG 4000 w/v, 0.2 M mannitol, 0.1 M CaCl_2_) for 10 min at room temperature. Transfection was stopped by adding double volume of W5 solution, followed by gentle centrifugation to pellet the protoplasts. The protoplasts were resuspended in W5 solution and incubated in the dark for 12 hr at plant growth room.

### Co-Immunoprecipitation (Co-IP) assays

The Co-IP assay was conducted with minor modifications to the protocol (*92*). Either 300 mg of ground transgenic Arabidopsis tissue from 6-day-old seedlings or protoplast samples were suspended in 1.0 mL of ice-cold Co-IP buffer (100 mM Tris-HCl, pH 7.5, 150 mM NaCl, 1 mM EDTA, 1 % Triton X-100, 3 mM DTT, 1 mM PMSF, 1 x protease inhibitor cocktail) in a 1.5 mL LoBind tube. The samples were centrifuged at 13000 rpm for 10 min at 4 °C to remove cell debris. A 45 µL aliquot of the supernatant was mixed with 15 µL of 4× Laemmli buffer and saved as the input sample. The remaining supernatant was transferred to a new 1.5 mL LoBind tube containing 20 µL of RFP-Trap magnetic agarose (ChromoTek, Cat # rtma) or GFP-Trap magnetic agarose (ChromoTek, Cat # gtma-100) and incubated on an end-to-end rotor at 4 °C for 1 hr. After incubation, the magnetic agarose beads were separated using a magnetic rack and washed three times with Co-IP buffer, followed by a final wash with 50 mM Tris-HCl (pH 8.0). The beads were resuspended in 30 µL of 50 mM Tris-HCl and mixed with 10 µL of 4 x Laemmli buffer. The mixture was boiled for 5 min and analyzed by immunoblotting.

### AlphaScreen-based protein-protein interaction assay

The AlphaScreen assay is a robust proximity-based approach to detect biomolecular interactions *in vitro* (*63*). It relies on the energy transfer between donor and acceptor beads brought into proximity by the binding of tagged proteins, resulting in a luminescent signal proportional to the interaction strength. The promoter, CDS, and terminator regions of the *PICALM-bls*, *AGIA-ERendo*, and *GFP-AGIA* constructs were amplified via PCR with SPu and AODA2306 primers and used as templates for *in vitro* transcription and translation in a wheat germ cell-free system (No Cat # CFS-TRI-1240, CellFree Sciences, Japan). Biotinylation at the biotin ligation site (bls) was performed enzymatically with BirA biotin ligase and 0.5 µM d-biotin (No Cat # 04822-04, Nacalai Tesque, Japan) as previously described(*93*). Protein synthesis was confirmed by immunoblotting with HRP-conjugated anti-AGIA (1:10,000) and anti-biotin (1:1,000, Cell Signaling Technology) antibodies. AlphaScreen reactions were performed with slight modifications to the protocol (*94*). A 15 μL reaction containing AlphaScreen buffer (100 mM Tris-HCl, pH 8.0, 0.1% Tween-20, 1 mg/mL BSA), 1 μL monobiotinylated PICALMs, and 1 μL AGIA-tagged ER-KD or GFP was incubated at 26°C for 1 hr in an Optiplate-384 plate (Revvity). A 10 μL detection mixture containing streptavidin-conjugated donor beads, protein A-conjugated acceptor beads, and anti-AGIA antibody in AlphaScreen buffer was then added. After an additional 1 hr incubation at 26°C, chemiluminescence signals were measured using an EnVision 2105 multimode plate reader (Revvity).

### Immunoblot analysis

Protein samples were resolved on 8 - 10% SDS-PAGE gels and transferred to PVDF membranes (BioRad Cat # 1620177). Membranes blocked in 5% nonfat milk in TBST for 1 hr at room temperature, followed by overnight incubation at 4 °C with primary antibodies, including anti-HA (1:5000, Abcam, Cat # Ab18181), anti-GFP (1:5000, Invitrogen, Cat # 33-2600), anti-RFP (1:2000, ChromoTek, Cat # pabr1), anti-FLAG (1:5000, Sigma, Cat # F3165) and anti-ACTIN (1:10,000, Abcam, Cat # Ab230169). After washing, membranes were incubated with HRP-conjugated secondary antibodies (1:10,000, Jackson ImmunoResearch Labs, Cat # 112-035-003 for rat, Cat # 115-035-062 for mouse) for 1 hr at room temperature. For PL samples, membranes were incubated with horseradish peroxidase conjugate streptavidin (1:5000, Invitrogen, Cat # S911). Protein bands were detected using chemiluminescence ECL substrate and imaged with a Bio-Rad ChemiDoc system.

### RT-qPCR analysis

Total RNA extraction, first-strand cDNA synthesis and RT-qPCR were performed as previously described(*77*). Gene expression levels were quantified using relative 2^−ΔCT^ method with the Arabidopsis *ACT2* (AT3G18780) as the endogenous reference for normalization. Primer sequences used were listed in Table S2.

### Quantification and statistical analysis

To quantitatively analyze the stomatal phenotypes of mutants and complementation lines, the abaxial side of Arabidopsis cotyledons was imaged using confocal microscopy. The images were used to classify and count different epidermal cell types. The stomatal index, defined as the number of stomata divided by the total number of epidermal cells, was calculated. The stomatal cluster index, representing the proportion of stomatal groups with two or more adjacent stomata, was calculated as the ratio of stomata in cluster groups to the total number of stomata. For each genotype, 8 - 20 cotyledons encompassing over 600-1500 epidermal cells were analyzed to ensure statistical robustness. To quantitatively analyze the AlphaScreen results, the mean chemiluminescence signals from three replicates were calculated and compared between ERECTA and GFP reactions. For mass spectrometry data, protein fold-changes were calculated as log2-transformed ratios of normalized PSMs between experimental and control conditions. Protein enrichment significance was assessed using a one-sided Z-test (*95*) with a 95% confidence threshold (z ≥ 1.645). *p*-values were calculated using the pnorm function in R (version 4.2.2) and adjusted for multiple comparisons using the Benjamini-Hochberg false discovery rate correction as previously described (*96*). Statistical analyses and data visualization were performed using R. For the two-sample comparison, either Student’s t-test or Welch’s t-test was performed. For the multiple sample comparison, one-way ANOVA with post-hoc Tukey HSD test was performed. The corresponding *p*-values and sample size (n) are provided in the respective figure legends.

## Acknowledgments

We thank Rachael Cox for developing the R package used in this study for differential protein interaction analysis. We also show our gratitude to Kaysa Kerr, Nicole Uwakwe, Geovanny Zarceno, and Calvin Coffin for assisting in the screening of transgenic and T-DNA insertion mutants, Katsunari Maruyama for assisting with *in vitro* interaction assays, and Krishna Sepuru and Liangliang Chen for insightful discussion.

## Funding

US National Institute of General Medical Sciences R35GM122480 (EMM)

US Army Research Office W911NF-12-1-0390 (EMM)

The Welch Foundation F-1515 (EMM)

MEXT Japan Society for the Promotion of Science KAKENHI JP24K02050 (TU)

The Takeda Science Foundation (TU)

Japan Science and Technology Agency CREST JPMJCR20E5 (KE)

Japan Science and Technology Agency ERATO (JPMJER2403) (KE)

Howard Hughes Medical Institute (KUT)

The start-up fund from The University of Texas at Austin (KUT)

The Johnson & Johnson Centennial Chair (KUT)

American Society of Plant Biologists Undergraduate Summer Research Fellowship (CK)

## Author contributions

Conceptualization: PFB,

Methodology: PFB, TU, EMM, KUT

Investigation: PFB, CK, MHV, KE, AN

Bioinformatics Analysis: PFB, OP, EMM

Visualization: PFB, EMM, KUT

Supervision: KUT, EMM, TU

Writing—original draft: PFB, KUT

Writing—review & editing: PFB, CK, TU, KE, OP, EMM, KUT

## Data and materials availability

The mass spectrometry data generated for the proteomic analysis in this study have been deposited to the ProteomeXchange consortium via the partner repository MassIVE (https://massive.ucsd.edu/ProteoSAFe/static/massive.jsp) with the dataset identifier: MSV000096929. Original western blot images have been deposited to Texas Dataverse (https://doi.org/10.18738/T8/RUXBAT). This paper does not report original code.

